# Efficient Coding of Spatial Frequency in Natural Images: Cross-frequency Dependence

**DOI:** 10.1101/2025.09.18.677139

**Authors:** Linshan Wang, Reza Farivar

## Abstract

Research suggests that spatial frequency (SF) channels in the visual system operate with a degree of independence. However, the independence model has been questioned by evidence of non-additive effects in compound gratings, indicating complex interactions between SF channels. These studies, however, typically employ artificial stimuli, leaving questions about SF processing in natural images. Efficient Coding hypothesis, which posits that the visual system minimizes redundancy and retains relevant information, predicts a dependence between HSF and LSF. In this study, we examined interactions between LSF and HSF using natural and phase-scrambled images to explore SF integration during perception. Participants completed an SF identification task, using both natural and scrambled images to isolate the role of phase alignment. Our results indicate that HSF and LSF interact primarily in phase-aligned conditions, with phase scrambling driving independent processing of two SFs and reducing error rates. These findings suggest that phase alignment enhances perceptual efficiency, facilitating a trade-off between accuracy and redundancy reduction in natural scene processing.

## 1. Introduction

Many studies have supported the notion that spatial frequency (SF) channels operate independently (Stecher et al., 1973; Stromeyer & Klein, 1974). Early electrophysiological evidence, such as recorded responses from individual neurons in cat primary visual cortex (V1), demonstrated that V1 neurons respond optimally to specific spatial frequencies (Maffei & Fiorentini, 1973). De Valois and colleagues extended these findings to primates, showing that macaque V1 neurons are similarly selective to certain SF (1982). The organization of SF tuned neurons in V1 suggests some degree of independence between SF channels as well, with weak clustering of neurons implying that local interactions are mainly limited to neurons with similar SF preferences, as seen in studies by Tolhurst and Thompson (1982) and DeAngelis et al. (1999).

Psychophysical studies have also supported the idea of independence between SF channels. Strong masking effects were found when the target and mask shared the same SF but minimal interference when they differed, indicating that SF channels primarily interact within their own frequency range (Legge & Foley, 1980). Using an adaptation paradigm, when observers are adapted to a specific SF, sensitivity to that frequency decreases while sensitivity to other SFs remains largely unaffected (Blakemore & Campbell, 1969).

However, some psychophysical evidence challenged this model of independent SF channels by demonstrating that adaptation to a compound grating resulted in less threshold elevation than adaptation to a single SF (Graham et al., 1978; Nachmias et al., 1973; Wandell, 1995). This suggests that the effects of adaptation from individual SF components cannot simply be summed to predict the effects of a mixture, indicating that SF channels interact in a more complex, non-additive way.

Generalizing from the above-mentioned studies may be difficult, as the bulk of them have used simple synthetic stimuli such as gratings, whereas we now know that since neural system evolve under natural environment (Niven & Laughlin, 2008; Simoncelli & Olshausen, 2001) the cortex responds differently to features within natural images (Coggan et al., 2017; Farishta et al., 2022; Felsen et al., 2005; Talebi & Baker, 2012), SFs in natural images are represented in far more complex ways than simple gratings. LSF provide a coarse, contextually rich background, while HSF add detailed edges and textures, creating a layered and contextually meaningful visual representation (Marr, 2010; Walther et al., 2020). This interaction allows the human visual system to rapidly interpret images holistically, blending broad contextual information with specific details in a way that simple gratings cannot achieve (Petras et al., 2021; Petras et al., 2019; Skyberg et al., 2022).

Haun and Peli (2013) tested how different SFs interaction contribute to perceived contrast of broadband natural images. Subjects were shown pairs of natural images with random variations in contrast amplitude at different SFs, and were asked to judge which image had higher overall contrast. Their results suggested that suppression is biased towards LSF, which they interpreted this result as energy efficiency—given that edges in images stimulate neural responses across multiple scales, the system may prioritize responses from HSF over LSF contrasts. This process may enhance efficiency by discarding information that can be recovered from HSF. However, their methods relied on the perception of contrast changes to infer potential interactions and did not explicitly manipulate SFs to directly observe how they interact.

Natural scene is highly redundant (Frazor & Geisler, 2006; Hoyer & Hyvärinen, 2000; Thomson, 1999). Different SF components provide similar visual experience at the same location in natural scene, termed as a ‘suspicious coincidence’, which is the consequence of Fourier phase-alignment of LSF and HSF (Zhaoping (2014), p185). Visual system should take advantage of the phase-alignment present in natural images to enhance perceptual efficiency. The Efficient Coding hypothesis provides a useful framework for understanding why SF channels should not function independently but rather interact to enhance perceptual efficiency, in response to natural stimuli. According to this principle, the brain optimizes the encoding of sensory information by reducing redundancy and preserving the most relevant information, all while minimizing energy consumption (Atick & Redlich, 1990; Attneave, 1954; Barlow, 1961; Niven & Laughlin, 2008).

We sought to understand the extent to which SF channels may interact to alter perception of one another—a strong form of dependence. Using a factorial combination of two levels of low and high SF bands in natural images as compared to a phase-scrambled control set of images, we measured participants’ identification performance and then tested the linearity vs. non-linearity of the interactions of the SF bands.

## 2. Materials and methods

### 2.1 Stimuli generation

Stimuli were generated from 33 object-based natural images retrieved from the THINGS database (Hebart et al., 2019). 3 were used in the training phase and 30 in the testing phase. To standardize presentation, all images were cropped to a square aspect ratio and resized to a resolution of 512 × 512 pixels. SF manipulations were applied directly to the colored images, rather than grayscale-converted versions. Specifically, filtering was performed separately on each RGB channel using the Difference of Gaussian (DoG) method to isolate low spatial frequency (LSF) and high spatial frequency (HSF) components. While many SF studies employ grayscale images to reflect luminance-based processing, we retained color information to preserve the ecological richness of natural image inputs. We acknowledge that cortical color processing does not map one-to-one onto RGB channels; however, our aim was to preserve naturalistic correlations between SF and chromatic content rather than to model isoluminant mechanisms. This choice is consistent with previous studies investigating SF interactions in colored natural images.

For each original image, we generated two levels of LSF-filtered and two levels of HSF-filtered versions. Final experimental stimuli were created by pairwise recombination of one LSF and one HSF level, yielding four SF combinations: HH, HL, LH, and LL. The naming convention follows the format level + SF, where level denotes high (h) or low (l) relative frequency content, and SF specifies the frequency band (LSF or HSF). For example, hHSF refers to the higher-frequency HSF component, and lLSF refers to the lower-frequency LSF component.

We deliberately chose to retain color rather than convert the images to grayscale, as grayscale conversion would have removed chromatic information that naturally co-varies with spatial frequency content, thereby reducing ecological validity. In natural environments, color and luminance are jointly structured across spatial scales (Simoncelli & Olshausen, 2001), and eliminating color collapses this relationship, disproportionately emphasizing luminance contrasts while discarding chromatic edges that support object and scene recognition. Prior work further shows that chromatic information can modulate spatial frequency tuning in both early and mid-level visual areas (Johnson et al., 2001; Mazer et al., 2002), and that color–luminance correlations are integral to redundancy reduction under efficient coding (Kay & Yeatman, 2017).

Each experimental stimulus consists of one LSF band with one HSF band. To derive these bandpass components, the DoG filtering technique was applied, which involved calculating the difference between two blurred versions of each image. The blurring was achieved using Gaussian kernels with different standard deviations (shown in Figure 1). The radii of the Gaussian kernels were determined by multiplying the standard deviation by four and rounding to the nearest integer. The image processing pipeline was carried out using Python 3.8, using OpenCV (Bradski, 2000) for image format conversion, SciPy (Virtanen et al., 2020) for Gaussian filtering, and NumPy (Harris et al., 2020) for numerical operations.

**Figure 1.**
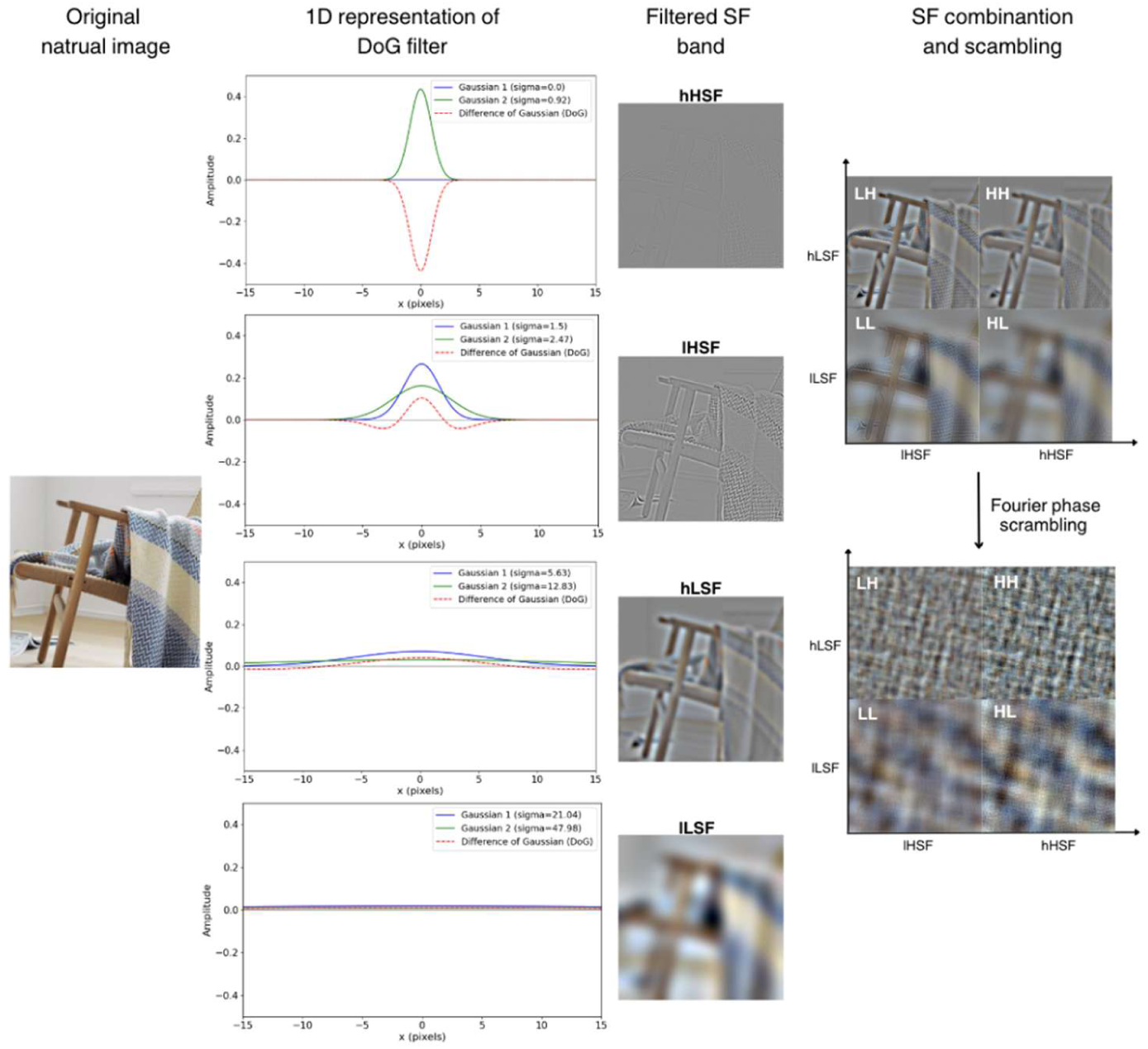
Stimulus preparation pipeline. An example natural image (left) was filtered into spatial frequency (SF) bands using the Difference-of-Gaussians (DoG) method. The second column shows 1D representations of the DoG filters used to extract high spatial frequency (HSF) and low spatial frequency (LSF) components at two levels (hHSF, lHSF, hLSF, lLSF). The third column illustrates example outputs of the filtering process for each band. These filtered components were then recombined to create four SF combinations: HH (hHSF + hLSF), HL (hHSF + lLSF), LH (lHSF + hLSF), and LL (lHSF + lLSF) (fourth column, top). To disrupt the natural phase alignment between frequency bands, Fourier phase scrambling was applied to the recombined stimuli (fourth column, bottom), yielding images with preserved amplitude spectra but no structure.

Due to the nature of the DoG filtering process, we could not set a precise cut off for each SF band. Despite this, DoG filter was chosen for its ability to provide smooth transition and mitigate potential ringing effects on filtered natural images, which can be difficult to achieve by other filtering methods such as log-Gabor and Heaviside (Walther et.al, 2020).

To estimate cycles per degree (cpd) from each band, a Fourier Transform is applied to the image set to convert it into the frequency domain, where the Power Spectral Density (PSD) is computed to quantify the energy distribution across spatial frequencies. After normalizing the PSD and removing the DC, the radial frequency, calculated as the Euclidean distance from the center, is converted to cycles per picture (cpp), this frequency is then converted to cpd by determining the visual angle subtended by a single pixel, calculated using the pixel size (0.0276cm) and the viewing distance (57 cm). We estimated the peak SF for each band: LSF ≈ 3.0 cpp = 0.21 cpd; hLSF ≈ 6.5 cpp = 0.46 cpd; lHSF ≈ 12.0 cpp = 0.85 cpd; hHSF ≈ 128.0 cpp = 9.05 cpd.

Phase scrambling was carried out in the Fourier domain separately for each RGB channel. For each channel, the image was decomposed into its amplitude spectrum (which reflects the distribution of spatial frequencies) and its phase spectrum (which encodes spatial structure). The original phase values were replaced with randomly assigned values drawn from a uniform distribution between −π and π, while the amplitude spectrum was preserved. Each scrambled channel was then reconstructed using the inverse Fourier transform, and the three scrambled channels (R, G, and B) were recombined to form the final scrambled image. This procedure disrupted spatial phase structure while preserving the overall amplitude spectrum and color balance of the original images.

RMS and intensity normalization was applied after the recombination of SF components, not at the level of the individual LSF- or HSF-filtered images. Each recombined stimulus was normalized using a z-score transformation followed by rescaling to fixed target mean luminance and RMS contrast values (127.0 and 25.0, respectively). Specifically, for each pixel intensity 𝐼_norm_ 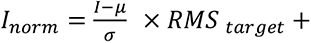 𝜇_target_, where 𝜇 and 𝜎 are the mean and standard deviation (RMS contrast) of the original recombined image, and 𝜇_target_ = 127.0 and 𝑅𝑀𝑆 _target_ = 25.0. This ensured that all final stimuli were equated in terms of global luminance and contrast energy. We confirmed this by logging the mean and RMS values for each image and verified that they were tightly controlled across conditions.

### 2.2 Participants

Fourteen participants (8 females) were recruited for the study. All provided informed consent prior to participation, and the experimental protocol approved by the Research Ethics Board of the McGill University Health Centre (MUHC), and all participants provided written informed consent prior to participation, in accordance with the Declaration of Helsinki. All participants reported normal or corrected-to-normal visual acuity and no history of neurological or visual disorders. Data from two participants was excluded due to inadequate performance (see below), leaving a final sample of twelve participants for analysis.

### 2.3 Procedure

There was no time limits imposed in any of the training or testing stages, and participants could take as much time as needed to make their decisions. There were no limits on the number of trials during the training phase, allowing participants to proceed until they met the accuracy requirements. Additionally, the stimuli used in training stage are not the same as in testing stage, to avoid practice effects.

The task was to identify the spatial frequency (SF) band combination of each stimulus in a four-alternative forced-choice (4AFC) design. The procedure was identical for natural and phase-scrambled conditions. Participants were seated in a dark room at a fixed viewing distance of 57 cm from the screen. The display had a mean luminance of 30.75 cd/m², a resolution of 1280 × 1024 pixels, and a refresh rate of 60 Hz. Gamma calibration was performed with a ColorCAL (Cambridge Research Systems) to ensure linear luminance output, and the experiment was programmed in PsychoPy (Peirce et al., 2019).

Participants were allowed to move their eyes freely during the task. To prevent reliance on global edges, a raised cosine window was applied to all images. There were no time limits in either training or testing, and participants could take as long as needed to make their responses. Importantly, the images used in training were different from those used in testing, ensuring that participants could not rely on practice effects.

To record their responses, participants pressed the keyboard key corresponding to the numerical type label assigned to each of the four possible SF combinations: type1 = HH (high LSF + high HSF), type2 = HL (high LSF + low HSF), type3 = LH (low LSF + high HSF), and type4 = LL (low LSF + low HSF).

### Training Phase

Training was divided into two stages.

#### 1. Pairwise Training (2AFC)

Participants first completed six pairwise discrimination sessions, each contrasting two of the four SF combinations. Each trial presented a single stimulus (one image with a particular SF combination) at the center of the screen, and participants judged which SF combination was presented.

Each session consisted of 12 consecutive trials (3 base images × 4 possible combinations). The criterion for passing was ≤1 error within a block of 12 trials. If more than one error occurred, the session repeated with a randomized trial order until criterion was reached.

#### 2. Full Combination Training (4AFC)

After completing the pairwise training, participants moved on to the 4AFC training where all four SF combinations were presented in one session. The difficulty increased and allowed participants to further learn the difference between the four combinations. Here, participants indicated the complete SF combination (HH, HL, LH, or LL) by pressing one of four designated keys. The same accuracy criterion applied (≤1 error per 12 consecutive trials).

The stopping rule ensured that all participants reached high accuracy on a small set of images so that task performance during the testing phase could not be attributed to insufficient familiarization with the stimulus categories. By requiring criterion-level performance prior to testing, we minimized the likelihood that errors in the main experiment reflected misunderstanding of the task rather than genuine perceptual limitations.

### Testing Phase

In the main experiment, participants viewed 30 natural images in which each processed into four SF combinations (HH, HL, LH, LL), yielding 120 unique stimuli. Each set was repeated three times, for a total of 360 trials per condition (natural vs. scrambled). Trials were evenly distributed across six sessions, each consisting of 60 trials. Each trial displayed a single stimulus centrally until response. The 4AFC task required identifying whether the image contained high vs. low LSF and high vs. low HSF, corresponding to HH, HL, LH, or LL. To prevent decisional bias, each session included an equal number of the four SF combinations (15 per type).

Participants received real-time counters on the screen. The counters were intended to provide participants with real-time counters to monitor their performance and avoid decisional bias: The top row of the display indicated how many responses participants had made so far for each SF combination type (e.g., HH, HL, LH, LL). This helped ensure that participants did not unintentionally favor one response category. The bottom row indicated how many total stimuli had been displayed up to that point in the session.

In the feedback display, the “type” number indicated the correct SF combination for the stimulus on that trial, as explained above (Figure 2).

**Figure 2.**
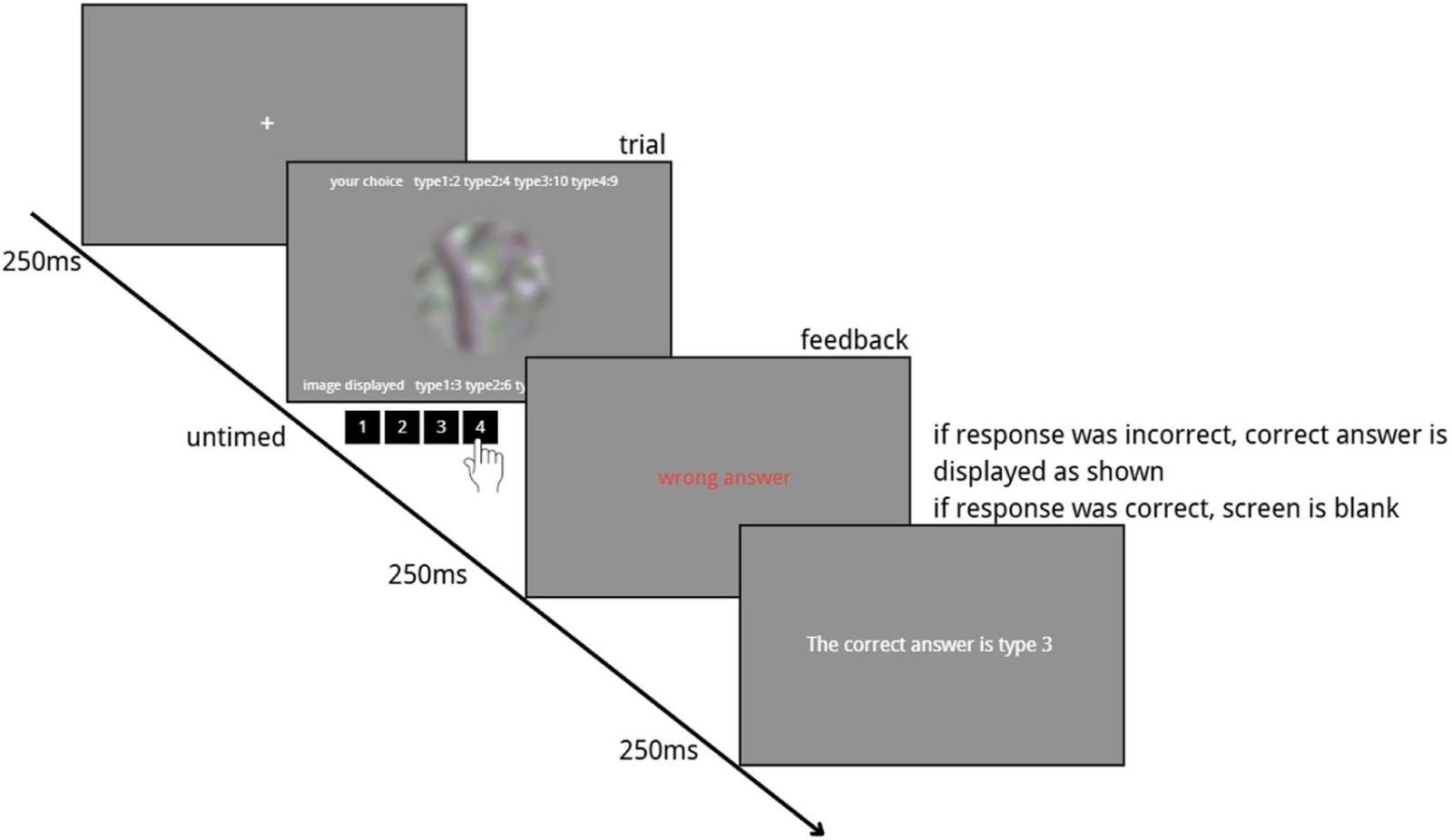
Schematic of the experimental procedure. Each trial consisted of a fixation period, the presentation of a single stimulus image and a 4AFC response screen where participants indicated the perceived SF combination. Only one image was shown per trial, not four simultaneously. The example trial sequence shown here is illustrative; the exact images and order of presentation were randomized across trials. The top row of counters displayed the number of responses made for each SF type to help participants avoid decisional bias, while the bottom row displayed the total number of stimuli presented. Feedback screens indicated the correct SF combination using the type code (e.g., type1 = HH, type2 = HL, type3 = LH, type4 = LL). Participants made responses using four designated keys on a standard keyboard, each consistently mapped to one SF combination.

## 3. Results

### 3.1. Accuracy of SF identification

It took on average about 30 minutes to learn the natural and 30 minutes for scrambled SF combinations. The SF identification task was challenging, as demonstrated by one participant’s inability to successfully pass the training phase in the natural image condition. As a result, we excluded the data from that participant. Of the 13 participants remaining, one participant excluded because his overall accuracy fell below 62.5%, the chance-level criterion for a 4AFC task. At the stimulus level, of the 30 raw images, 4 were excluded because their mean accuracy across participants was below 33%. When an image was removed, all four of its spatial-frequency variants (HH, LL, HL, and LH) were excluded together, to avoid unbalanced representation of conditions. These exclusion criteria ensured that the final dataset contained only reliable participants and stimuli, while maintaining comparability across all spatial-frequency combinations.

For the remaining stimuli, we computed mean accuracy for each unique stimulus by averaging across its three repetitions. This produced one accuracy score per stimulus per participant (26 images × 4 SF combinations × 2 conditions = 208 data points per participant). These values were then grouped according to image type (natural vs. scrambled), HSF (high vs. low), and LSF (high vs. low). A three-way repeated-measures ANOVA was conducted on these stimulus-level means to examine the main effects and interactions of image type, HSF, and LSF. There was a significant main effect of Condition, F(1, 200) = 22.28, p < .001, η²p = .10, indicating a large difference in accuracy between natural and scrambled images. The main effect of LSF approached significance, F(1, 200) = 3.79, p = .053, η²p = .02, suggesting a small influence of low spatial frequency. No main effect of HSF was found, F(1, 200) = 0.01, p = .943, η²p < .001.

The analysis revealed a significant Condition × LSF interaction, F(1, 200) = 4.67, p = .032, η²p = .02, a strong HSF × LSF interaction, F(1, 200) = 42.53, p < .001, η²p = .18, and a significant three-way Condition × HSF × LSF interaction, F(1, 200) = 9.38, p = .003, η²p = .04. The Condition × HSF interaction was not significant, F(1, 200) = 0.63, p = .429, η²p = .01.

To unpack the three-way interaction, follow-up ANOVAs were conducted separately for natural and scrambled conditions. In the natural condition, there was no effect of HSF, F(1, 100) = 0.40, p = .531, η²p < .01, but a significant main effect of LSF, F(1, 100) = 8.93, p = .004, η²p = .08 . Most importantly, the HSF × LSF interaction was highly significant, F(1, 100) = 48.60, p < .001, η²p = .33, representing a very large effect. In the scrambled condition, there were no significant main effects of HSF or LSF, Fs < 0.25, ps > .62, but the HSF × LSF interaction was significant, F(1, 100) = 5.67, p = .019, η²p = .05.

### 3.2 Confusion Pattern of SF identification

To further visualize participants’ performance, confusion matrices in Table 1 and 2 shows the mean percentage of stimulus-response mappings across the different SF combinations.

**Table 1.**
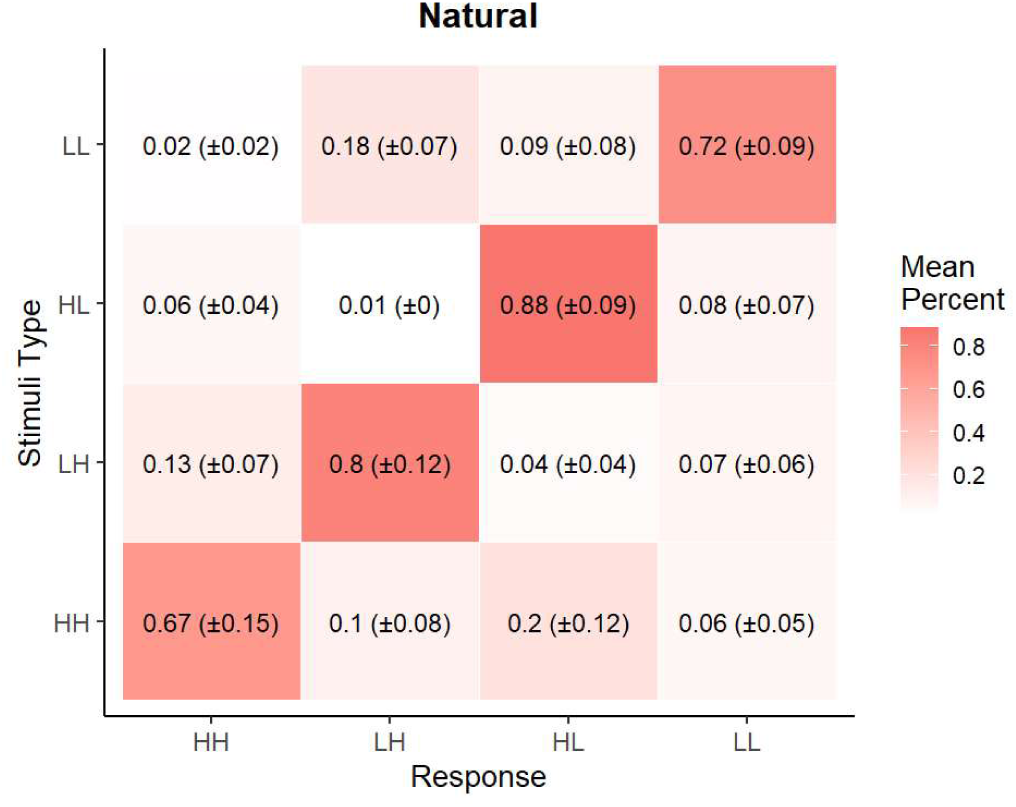
Confusion matrix in Natural condition. Mean response percentage on each SF combination, correct responses are shown along the diagonal.

**Table 2.**
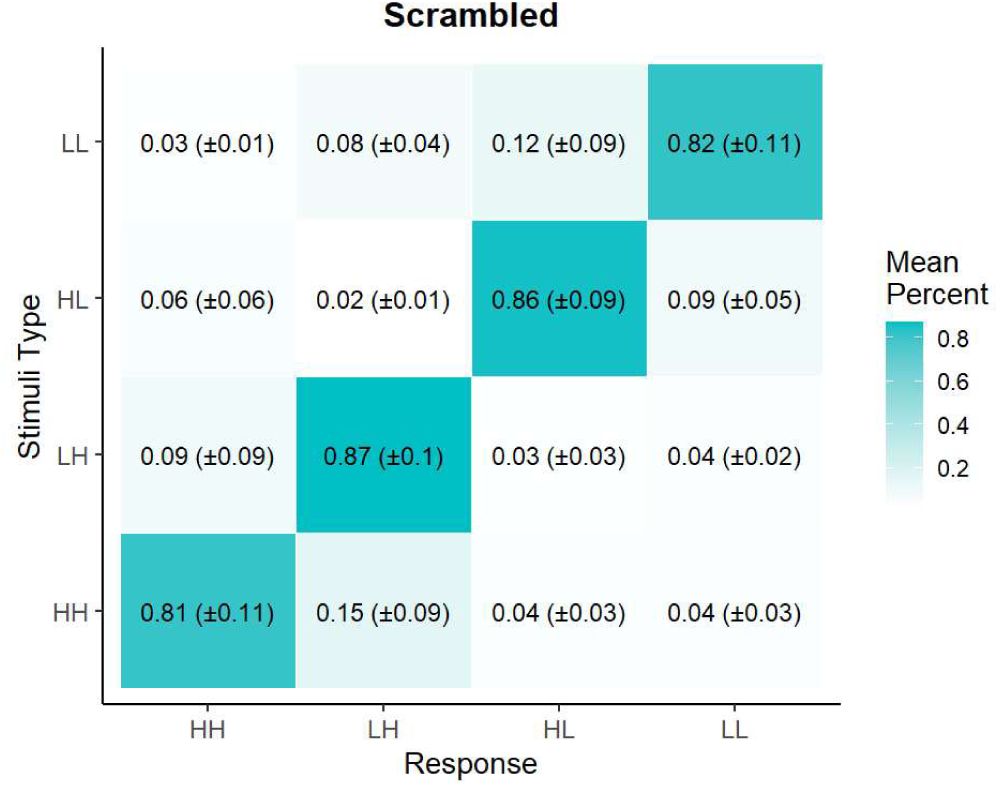
Confusion matrix in Scramble condition. Mean response percentage on each SF combination, correct responses are shown along the diagonal.

To better understand the nature of participants’ misclassifications, we categorized each incorrect response into one of five error types as shown in Figure 4. A two-way repeated-measures ANOVA was conducted on error proportions with error type (HSF at hLSF, HSF at lLSF, LSF at hHSF, LSF at lHSF, Both) and condition (natural vs. scrambled) as factors. The analysis revealed a significant main effect of error type, F(4, 238) = 19.69, p < .001, η²p = .25, indicating that the frequency of errors varied systematically across error categories. There was no significant main effect of condition, F(1, 238) = 0.05, p = .821, η²p < .001. Importantly, the interaction between error type and condition was significant, F(4, 238) = 14.36, p < .001, η²p = .19, showing that the relative distribution of error types differed between natural and scrambled images. Post hoc comparisons (Bonferroni-corrected) tested for differences between natural and scrambled conditions within each error type. There was no difference in the rate of “Both” errors between conditions, t(238) = 0.38, p = .702. In contrast, participants made significantly fewer HSF errors in the natural condition compared to scrambled, both at high LSF (t(238) = –4.09, p < .001, estimate = –0.0054) and at low LSF (t(238) = –3.89, p < .001, estimate = –0.0052). Conversely, participants made significantly more LSF errors in the natural condition compared to scrambled, both at high HSF (t(238) = 3.43, p < .001, estimate = 0.0045) and at low HSF (t(238) = 3.71, p < .001, estimate = 0.0049).

**Figure 3.**
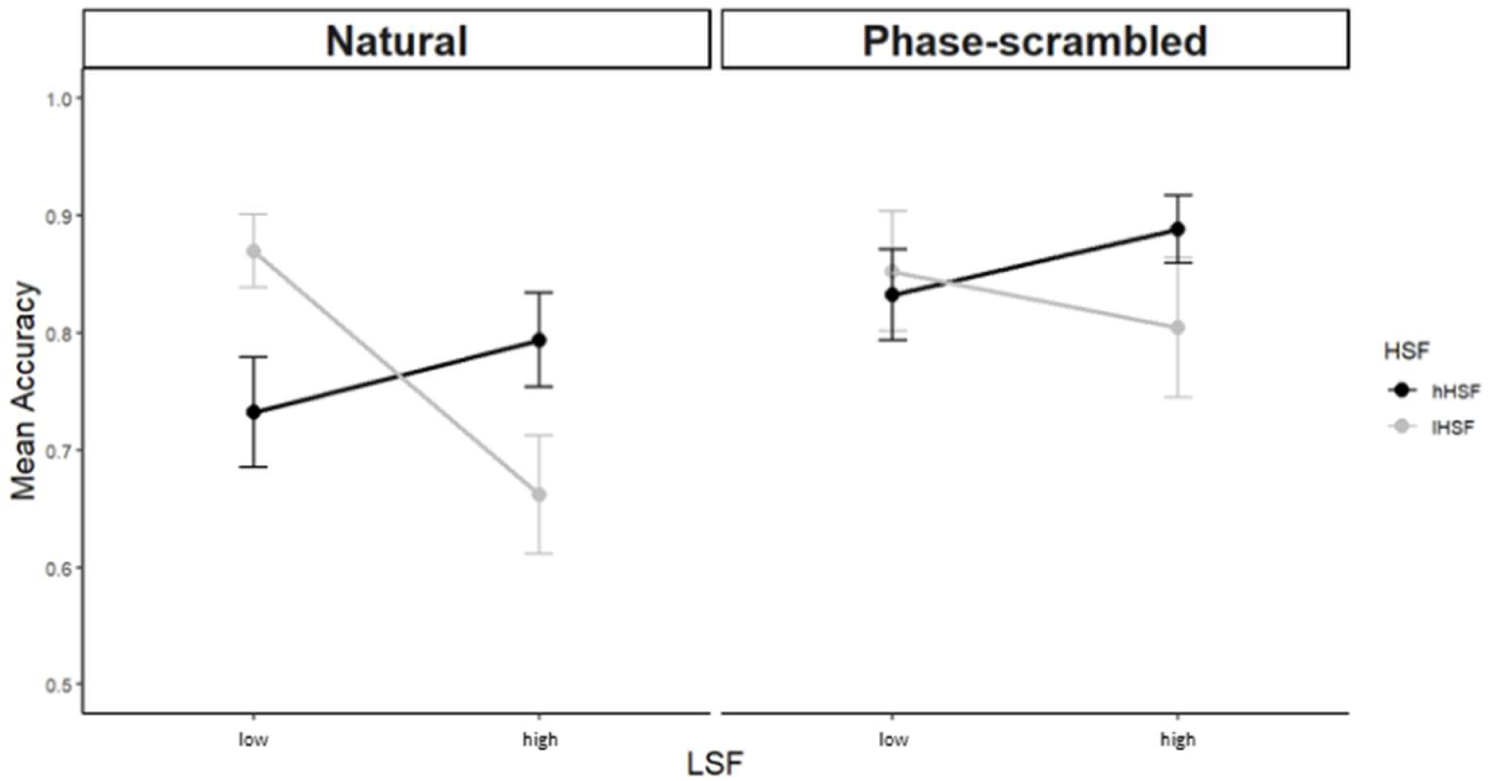
Mean accuracy for SF discrimination as a function of SF level and image condition. The x-axis shows the LSF level (low vs. high), while the y-axis shows participants’ mean accuracy (proportion correct) in identifying the SF combination. Results are plotted separately for natural images (left panel) and phase-scrambled images (right panel). Black lines represent high hHSF conditionsnand gray lines represent lHSF components,

**Figure 4.**
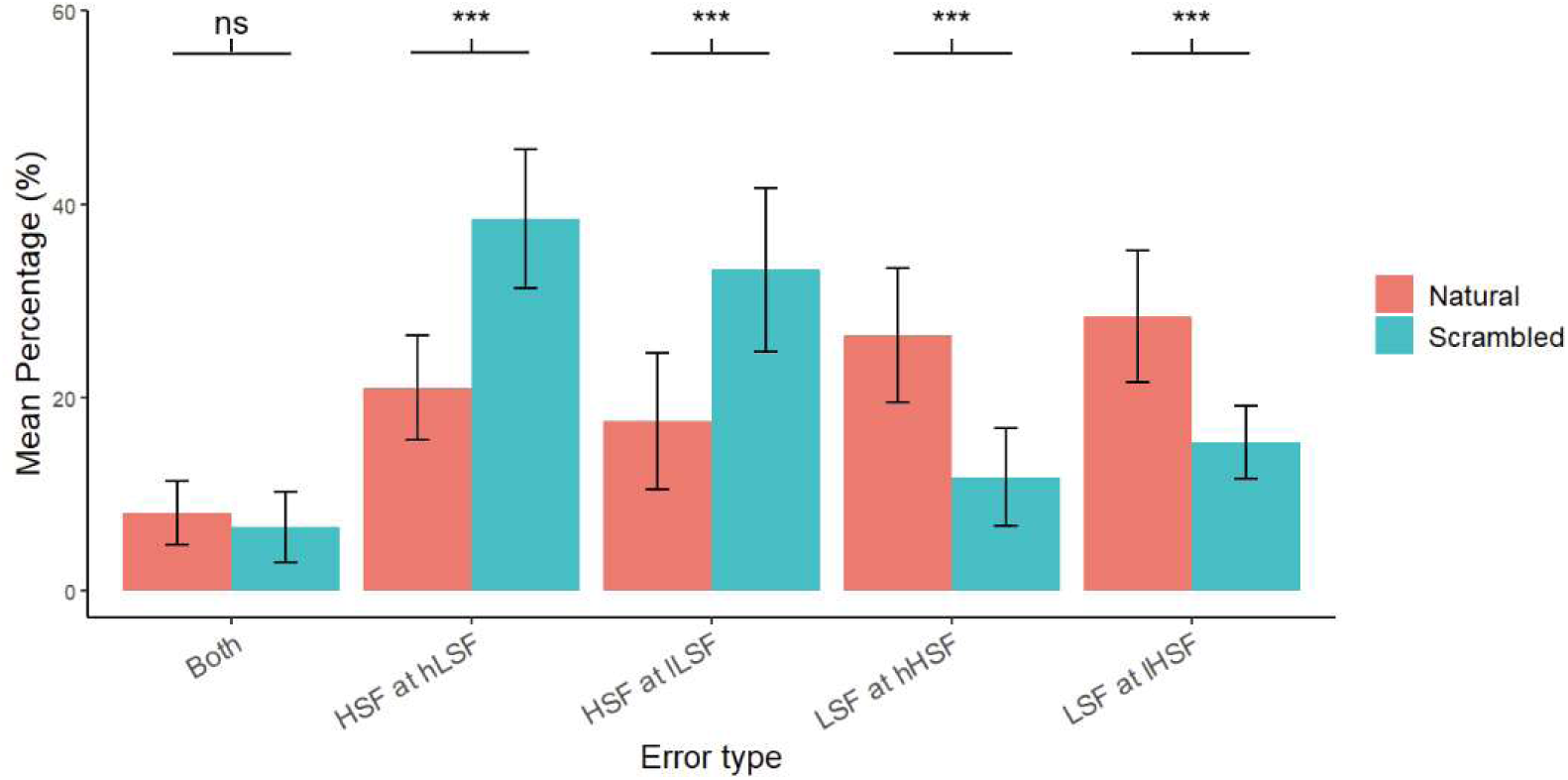
Error profiles across SF conditions in natural and scrambled images. Mean percentage of five error types is shown separately for natural and scrambled conditions: (1) Both – both HSF and LSF components misperceived; (2) HSF at hLSF – the high-frequency (HSF) component was reported incorrectly while the low-frequency (LSF) component was high; (3) HSF at lLSF – the HSF component was misperceived when the LSF component was low; (4) LSF at hHSF – the LSF component was misperceived when the HSF component was high; and (5) LSF at lHSF – the LSF component was misperceived when the HSF component was low. Error bars represent 95% confidence intervals.

## 4. Discussion

We applied a novel SF combination identification task to measure HSF and LSF perception when both components are present. Importantly, we tested this identification performance in natural and phase-scrambled conditions, which allowed us to make inferences about the potential for joint encoding of SF content in natural images. We showed that interaction between two channels is primarily driven by Fourier-phase alignment, which can be predicted by the Efficient Coding hypothesis, as this interaction was only observed on natural scene images.

Our results indicate that the perceptual effect of one SF component depends on the characteristics of the other. In other words, the influence of HSF on perception varies with the LSF component, and vice versa, suggesting that these two frequency channels do not operate entirely independently in visual processing. This finding aligns with the idea of prior psychophysical studies, emphasizing that combining simple sine gratings does not produce a purely additive effect, challenging earlier views that SF channels function independently.

The phase spectrum of natural images holds a significant portion of the perceptually important information, surpassing the amplitude spectrum in this regard (Oppenheim & Lim, 1981). In the Fourier domain, the phase spectrum holds positional information of SFs. When the phase is scrambled, the precise arrangement of structures in the image is disrupted. In other words, scrambling Fourier phases decorrelates HSF and LSF in terms of the visual information provided.

Our result suggested that in scrambled images, where phase relationships between HSF and LSF components are disrupted, the channels seem to operate more independently. This independence may simplify the processing demands on the visual system, which could account for the reduced error rates in this condition. Since phase alignment is a key feature of natural images, scrambling the Fourier phase of the images eliminates interactions between SF channels, allowing for more veridical perception.

This suggests a trade-off in visual processing—while the brain prioritizes efficiency by minimizing redundancy, this efficiency may come at the expense of precision. Redundancy reduction is essential for efficient coding, yet it can sometimes introduce perceptual errors. Previous research on V1 and MT neuron modelling demonstrated that illusions of direction can arise from mechanisms that reduce redundancy (Perrone & Liston, 2015). The different performance levels we observed between natural and phase-scrambled conditions highlight this trade-off between perceptual accuracy and redundancy reduction, as originally posited by Efficient coding hypothesis. Redundancy in vision is not inherently detrimental; rather, it facilitates error correction during information transmission when visual input is independent (Cortes et al., 2012; Zhaoping, 2014).

Natural images have a 1/f spatial frequency spectrum, with LSF containing more physical energy and HSF contributing less (Field, 1987; Ruderman & Bialek, 1994). If the brain merely reflected these raw statistics, we would expect errors to track energy, with more reliable LSF perception. Instead, our error profile analysis shows the opposite under natural conditions: errors are disproportionately weighted toward LSF, while HSF errors are minimized. This indicates a non-uniform coding strategy, where the brain strategically up-weights HSF information and suppresses LSF contributions when phase alignment allows reliable integration across scales. Such a strategy is consistent with principles of efficient coding: rather than redundantly encoding coarse features that dominate the stimulus energy, the visual system emphasizes the sparser, but more fine-scale edges. These findings align with Haun and Peli (2013) hypothesis that the visual system prioritizes edge details over coarse features in natural images, given the redundancy of LSF when edges are present. The increased LSF perceptual errors in natural images imply that, when SF channels function dependently, the brain selectively suppresses LSF information, likely to support efficient perception. Specifically, when phase alignment ensures that HSF and LSF components are well-integrated, the redundant coarse information provided by LSF becomes less critical, allowing the brain to emphasize the more informative HSF features.

These findings highlight a non-uniform sensitivity to phase alignment between HSF and LSF channels: HSF perception is critically dependent on phase alignment, whereas LSF perception is less so. This asymmetry aligns with Farivar et al. (2017), who demonstrated neural sensitivity of HSF to phase coherence, and extends it by showing how phase alignment determines the allocation of perceptual errors across frequency channels. More broadly, the results illustrate how the brain leverages efficient coding principles, suppressing redundant LSF signals in natural conditions to prioritize HSF edges, but reversing this weighting when phase coherence is lost.

It is worth considering whether our normalization procedure may have influenced the relative perceptual weighting of LSF and HSF information. Because normalization in our study was applied after recombination of the SF components, rather than to the individual LSF- and HSF-filtered images, the contribution of HSF may have been perceptually attenuated in certain conditions. At the extreme, if this bias led observers to effectively disregard HSF, we would expect HSF errors to dominate across all conditions. However, this was not observed: in natural images, errors were weighted toward LSF, indicating that the brain prioritized HSF features despite their lower physical energy. Only under phase scrambling did errors shift toward HSF, consistent with the view that HSF perception critically depends on phase alignment. Thus, the observed error reversal reflects a genuine phase-driven effect, not merely an artifact of stimulus normalization.

Our results demonstrate significant perceptual interactions between HSF and LSF in natural images but not in scrambled images, consistent with the Efficient Coding hypothesis. This framework posits that the brain optimizes sensory processing by maximizing the informational efficiency of neural representations. **Entropy** is used to quantify uncertainty in sensory input 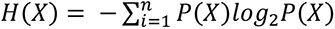, where 𝐻(𝑋) is used represent represents the amount of missing information or "surprise" before observing a specific input 𝑋. In natural images, the spatial alignment of features across SF bands creates interdependence between HSF and LSF, such that the probability of one SF band depends on the other:

This dependence reduces the **conditional entropy**, as knowing one SF component reduces uncertainty about the other: 𝐻(𝑆_high_l𝑆_low_) < 𝐻 (𝑆_high_) and 𝐻(𝑆_low_l𝑆_high_) < 𝐻 (𝑆_low_). Based on the formula of **mutual information** 𝐼(𝑋; 𝑌) = 𝐻(𝑋) − 𝐻(𝑋|𝑌), mutual information between the two components is calculated as 𝐼 (𝑆_high_; 𝑆_low_) = 𝐼 (𝑆_low_; 𝑆_high_) > 0 . This indicates that knowing one SF band provides significant information about the other, facilitating efficient encoding and interaction in natural images.

By contrast, in scrambled images, phase relationships across SF bands are disrupted, breaking the correlations between HSF and LSF. As a result, the two SF components are statistically independent. Consequently, 𝐻(𝑆_high_l𝑆_low_) = 𝐻 (𝑆_hig_) and 𝐻(𝑆_low_l𝑆_hig_) = 𝐻 (𝑆_low_), and mutual information vanishes:𝐼 (𝑆_high_; 𝑆_low_) =𝐼 (𝑆_low_; 𝑆_high_) = 0. This lack of shared information explains why no perceptual interaction is observed in scrambled images. Our results suggest that mutual information between SF bands in natural images allows the visual system to optimize sensory coding by reducing redundancy and uncertainty, enhancing perceptual efficiency.

An important consideration is the relative role of semantic versus structural factors in driving the observed LSF–HSF interactions. The dependencies we observed may partly reflect the semantic content of natural object images rather than purely statistical correlations, since natural images carry meaning and recognizability cues that could influence how observers integrate spatial frequency information. Indeed, prior work showed that phase disruption eliminates semantic structure and impairs scene understanding even when amplitude spectra are preserved (Hansen & Loschky, 2013);. At the same time, even in the absence of meaning, the visual system is sensitive to statistical properties of textures and synthetic images. Evidence from both neural and perceptual studies shows that responses to statistical image structure occur independently of semantic recognizability (Freeman & Simoncelli, 2011; Ziemba et al., 2016). This implies that statistical correlations alone are sufficient to drive responses. Thus, while our findings are consistent with the Efficient Coding Hypothesis, we cannot rule out an additional role of semantic content in shaping SF interactions. Future studies employing artificially generated, phase-aligned but semantically meaningless stimuli could help disentangle these contributions by testing whether the LSF–HSF interaction persists in the absence of recognizability.

## Reference

1. Atick, J. J., & Redlich, A. N. (1990). Towards a Theory of Early Visual Processing. Neural Computation, 2(3), 308–320. 10.1162/neco.1990.2.3.308

2. Attneave, F. (1954). Some informational aspects of visual perception. Psychological Review, 61(3), 183–193. 10.1037/h0054663

3. Barlow, H. (1961). Possible Principles Underlying the Transformations of Sensory Messages. Sensory Communication, 1. 10.7551/mitpress/9780262518420.003.0013

4. Blakemore, C., & Campbell, F. W. (1969). On the existence of neurones in the human visual system selectively sensitive to the orientation and size of retinal images. The Journal of Physiology, 203(1), 237–260. 10.1113/jphysiol1969.sp008862

5. Bradski, G. (2000). *The Opencv Library* (Vol. 25).

6. Coggan, D. D., Allen, L. A., Farrar, O. R. H., Gouws, A. D., Morland, A. B., Baker, D. H., & Andrews, T. J. (2017). Differences in selectivity to natural images in early visual areas (V1–V3. Scientific Reports, 7(1), 2444. 10.1038/s41598-017-02569-4

7. Cortes, J. M., Marinazzo, D., Series, P., Oram, M. W., Sejnowski, T. J., & Rossum, M. C. W. (2012). The effect of neural adaptation on population coding accuracy. Journal of Computational Neuroscience, 32(3), 387–402. 10.1007/s10827-011-0358-4

8. De Valois, R. L., Albrecht, D. G., & Thorell, L. G. (1982). Spatial frequency selectivity of cells in macaque visual cortex. Vision Res, 22(5), 545–559. 10.1016/0042-6989(82)90113-4

9. DeAngelis, G. C., Ghose, G. M., Ohzawa, I., & Freeman, R. D. (1999). Functional Micro-Organization of Primary Visual Cortex: Receptive Field Analysis of Nearby Neurons. The Journal of Neuroscience, 19(10), 4046. 10.1523/JNEUROSCI.19-10-04046.1999

10. Farishta, R. A., Yang, C. L., & Farivar, R. (2022). Blur Representation in the Amblyopic Visual System Using Natural and Synthetic Images. Investigative Opthalmology & Visual Science, 63(1), 3. 10.1167/iovs.63.1.3

11. Farivar, R., Clavagnier, S., Hansen, B. C., Thompson, B., & Hess, R. F. (2017). Non-uniform phase sensitivity in spatial frequency maps of the human visual cortex. The Journal of Physiology, 595(4), 1351–1363. 10.1113/JP273206

12. Felsen, G., Touryan, J., Han, F., & Dan, Y. (2005). Cortical Sensitivity to Visual Features in Natural Images. PLoS Biology, 3(10), 342– 342. 10.1371/journal.pbio.0030342

13. Field, D. J. (1987). Relations between the statistics of natural images and the response properties of cortical cells. J Opt Soc Am A, 4(12), 2379–2394. 10.1364/josaa.4.002379

14. Frazor, R. A., & Geisler, W. S. (2006). Local luminance and contrast in natural images. Vision Research, 46(10), 1585–1598. 10.1016/j.visres.2005.06.038

15. Freeman, J., & Simoncelli, E. (2011). Metamers of the ventral stream. Nature Neuroscience, 14(9), 1195–1201. 10.1038/nn.2889

16. Graham, N., Robson, J., & Nachmias, J. (1978). Grating summation in fovea and periphery. Vision Research, 18(7), 815–825.

17. Hansen, B. C., & Loschky, L. C. (2013). The contribution of amplitude and phase spectra-defined scene statistics to the masking of rapid scene categorization. Journal of Vision, 13(13), 21–21. 10.1167/13.13.21

18. Harris, C. R., Millman, K. J., Walt, S. J., Gommers, R., Virtanen, P., Cournapeau, D., Wieser, E., Taylor, J., Berg, S., Smith, N. J., Kern, R., Picus, M., Hoyer, S., Kerkwijk, M. H., Brett, M., Haldane, A., Del Río, J. F., Wiebe, M., Peterson, P., & Oliphant, T. E. (2020). Array programming with NumPy. Nature, 585(7825), 357–362. 10.1038/s41586-020-2649-2

19. Haun, A. M., & Peli, E. (2013). Perceived contrast in complex images. Journal of Vision, 13(13), 3–3. 10.1167/13.13.3

20. Hebart, M. N., Dickter, A. H., Kidder, A., Kwok, W. Y., Corriveau, A., Wicklin, C., & Baker, C. I. (2019). THINGS: A database of 1,854 object concepts and more than 26,000 naturalistic object images. PLOS ONE, 14(10), 0223792. 10.1371/journal.pone.0223792

21. Hoyer, P. O., & Hyvärinen, A. (2000). Independent component analysis applied to feature extraction from colour and stereo images. Network: Computation in Neural Systems, 11(3), 191–210. 10.1088/0954-898X_11_3_302

22. Johnson, E. N., Hawken, M. J., & Shapley, R. (2001). The spatial transformation of color in the primary visual cortex of the macaque monkey. Nature Neuroscience, 4(4), 409–416. 10.1038/86061

23. Kay, K. N., & Yeatman, J. D. (2017). Bottom-up and top-down computations in word- and face-selective cortex. eLife, 6, 22341. 10.7554/eLife.22341

24. Legge, G. E., & Foley, J. M. (1980). Contrast masking in human vision. Journal of the Optical Society of America, 70(12), 1458. 10.1364/JOSA.70.001458

25. Maffei, L., & Fiorentini, A. (1973). The visual cortex as a spatial frequency analyser. Vision Res, 13(7), 1255–1267. 10.1016/0042-6989(73)90201-0

26. Marr, D. (2010). Vision: A Computational Investigation into the Human Representation and Processing of Visual Information. The MIT Press. 10.7551/mitpress/9780262514620.001.0001

27. Mazer, J. A., Vinje, W. E., McDermott, J., Schiller, P. H., & Gallant, J. L. (2002). Spatial frequency and orientation tuning dynamics in area V1. Proceedings of the National Academy of Sciences of the United States of America, 99(3), 1645–1650. 10.1073/pnas.0226384999

28. Nachmias, J., Sansbury, R., Vassilev, A., & Weber, A. (1973). Adaptation to square-wave gratings: In search of the elusive third harmonic. Vision Research, 13(7), 1335–1342. 10.1016/0042-6989(73)90209-5

29. Niven, J. E., & Laughlin, S. B. (2008). Energy limitation as a selective pressure on the evolution of sensory systems. Journal of Experimental Biology, 211(11), 1792– 1804. 10.1242/jeb.017574

30. Oppenheim, A. V., & Lim, J. S. (1981). The importance of phase in signals. Proceedings of the IEEE, 69(5), 529–541. 10.1109/PROC.1981.12022

31. Peirce, J., Gray, J. R., Simpson, S., MacAskill, M., Höchenberger, R., Sogo, H., Kastman, E., & Lindeløv, J. K. (2019). PsychoPy2: Experiments in behavior made easy. Behav Res Methods, 51(1), 195–203. 10.3758/s13428-018-01193-y

32. Petras, K., Ten Oever, S., Dalal, S. S., & Goffaux, V. (2021). Information redundancy across spatial scales modulates early visual cortical processing. NeuroImage, 244, 118613. 10.1016/j.neuroimage.2021.118613

33. Petras, K., Ten Oever, S., Jacobs, C., & Goffaux, V. (2019). Coarse-to-fine information integration in human vision. NeuroImage, 186, 103–112. 10.1016/j.neuroimage.2018.10.086

34. Ruderman, D. L., & Bialek, W. (1994). Statistics of natural images: Scaling in the woods. Physical Review Letters, 73(6), 814–817. 10.1103/PhysRevLett.73.814

35. Simoncelli, E. P., & Olshausen, B. A. (2001). Natural Image Statistics and Neural Representation. Annual Review of Neuroscience, 24(1), 1193–1216. 10.1146/annurev.neuro.24.1.1193

36. Skyberg, R., Tanabe, S., Chen, H., & Cang, J. (2022). Coarse-to-fine processing drives the efficient coding of natural images in mouse visual cortex. Cell Reports, 38(13), 110606. 10.1016/j.celrep.2022.110606

37. Stecher, S., Sigel, C., & Lange, R. V. (1973). Spatial frequency channels in human vision and the threshold for adaptation. Vision Research, 13(9), 1691–1700. 10.1016/0042-6989(73)90088-6

38. Stromeyer, C. F., 3rd, & Klein, S. (1974). Spatial frequency channels in human vision as asymmetric (edge) mechanisms. Vision Res, 14(12), 1409–1420. 10.1016/0042-6989(74)90016-9

39. Talebi, V., & Baker, C. L., Jr. (2012). Natural versus synthetic stimuli for estimating receptive field models: a comparison of predictive robustness. J Neurosci, 32(5), 1560–1576. 10.1523/jneurosci.4661-12.2012

40. Thomson, M. G. A. (1999). Higher-order structure in natural images. Journal of the Optical Society of America A, 16(7), 1549. 10.1364/JOSAA.16.001549

41. Tolhurst, D. J., & Thompson, I. D. (1982). Organization of neurones preferring similar spatial frequencies in cat striate cortex. Exp Brain Res, 48(2), 217–227. 10.1007/bf00237217

42. Virtanen, P., Gommers, R., Oliphant, T. E., Haberland, M., Reddy, T., Cournapeau, D., Burovski, E., Peterson, P., Weckesser, W., Bright, J., Walt, S. J., Brett, M., Wilson, J., Millman, K. J., Mayorov, N., Nelson, A. R. J., Jones, E., Kern, R., Larson, E., & Vázquez-Baeza, Y. (2020). SciPy 1.0: Fundamental algorithms for scientific computing in Python. Nature Methods, 17(3), 261–272. 10.1038/s41592-019-0686-2

43. Walther, D. B., Perfetto, S., & Wilder, J. (2020). Spatial Frequency Filtering: Choices Matter. Journal of Vision, 20(11), 1205–1205. 10.1167/jov.20.11.1205

44. Wandell, B. A. (1995). Foundations of vision. Sinauer Associates.

45. Zhaoping, L. (2014). Understanding Vision: Theory, Models, and Data (1st ed.). Oxford University PressOxford. 10.1093/acprof:oso/9780199564668.001.0001

46. Ziemba, C. M., Freeman, J., Movshon, J. A., & Simoncelli, E. P. (2016). Selectivity and tolerance for visual texture in macaque V2. Proceedings of the National Academy of Sciences of the United States of America, 113(22), 3140–3149. 10.1073/pnas.1510847113

